# Multi-Tissue Multi-Compartment models of diffusion MRI

**DOI:** 10.1101/2021.01.29.428843

**Authors:** Matteo Frigo, Rutger H.J. Fick, Mauro Zucchelli, Samuel Deslauriers-Gauthier, Rachid Deriche

## Abstract

State-of-the-art multi-compartment microstructural models of diffusion MRI (dMRI) in the human brain have limited capability to model multiple tissues at the same time. In particular, the available techniques that allow this multi-tissue modelling are based on multi-TE acquisitions. In this work we propose a novel multi-tissue formulation of classical multi-compartment models that relies on more common single-TE acquisitions and can be employed in the analysis of previously acquired datasets. We show how modelling multiple tissues provides a new interpretation of the concepts of signal fraction and volume fraction in the context of multi-compartment modelling. The software that allows to inspect single-TE diffusion MRI data with multi-tissue multi-compartment models is included in the publicly available Dmipy Python package.

## 1. Introduction

Diffusion MRI (dMRI) is an imaging technique that allows to inspect the brain tissue microstructure in-vivo non-invasively. One of the most commonly studied microstructural feature is the *volume fraction* of a tissue in a sample. In particular, the intra-axonal (a.k.a. intra-cellular - IC), extra-axonal (a.k.a. extra-cellular - EC) and isotropic (ISO) or cerebro-spinal fluid (CSF) volume fractions have been investigated in the past literature with several models. Among all, we mention the neurite orientation dispersion and density imaging (NODDI) (Zhang et al., 2012) and NODDI-X (Farooq et al., 2016), ActiveAx (Alexander et al., 2010), the multi-compartment microscopic diffusion imaging framework (Kaden et al., 2016), the CHARMED model (Assaf and Basser, 2005), the intravoxel incoherent motion model (Le Bihan et al., 1988), the Stanisz model (Stanisz et al., 1997), the AxCaliber model (Assaf et al., 2008), the ball and stick model (Behrens et al., 2003), the Bingham-NODDI model (Tariq et al., 2016), FERNET (Parker et al., 2020), CODIVIDE (Lampinen et al., 2017), COMMIT (Daducci et al., 2015), VERDICT (Panagiotaki et al., 2014) and the DIAMOND model (Scherrer et al., 2016). The differences between these models lie on the representation employed in describing the tissue-specific signal and on the assumptions made on the model parameters. For example, intra-axonal diffusion can be modelled as the diffusion within a stick or a cylinder and some models fix the value of the diffusivity or tortuosity. A unifying aspect that characterizes most of the brain microstructure models is the *building-blocks* concept behind their formalisation. In other words, models are defined in a multi-compartment (MC) fashion, where the dMRI signal is described as a linear combination of single-tissue models. The resulting models are called MC models and they require the acquisition of multi-shell dMRI data in order to accurately disentangle the contribution of each compartment (Scherrer and Warfield, 2010). Thorough reviews have been dedicated to the design and validation of such models (Jelescu and Budde, 2017), to the sensitivity of MC models to experimental factors and microstructural properties of the described tissues (Afzali et al., 2020), and to the abstraction of these models that allows to obtain a unified theory (Fick et al., 2019).

Recent studies have highlighted that all of the available MC models are transparent to the *T*_2_ relaxation times of the modelled tissues (Veraart et al., 2018; Lampinen et al., 2019). As a consequence, they implicitly assume that all the considered tissues have the same non-diffusion weighted signal *S*_0_. While this is a reasonable assumption in some contexts, it is not true in general. In fact, each brain tissue is characterized by a specific relaxation time which makes *T*_2_ imaging possible. Assuming that all the tissues have a single *S*_0_ response simplifies the model at the cost of biophysical accuracy. Tissue fractions obtained with this assumption are called *signal* fractions, in contrast with the unbiased *volume* fractions which can be obtained with models that account for different *S*_0_ responses of the modelled tissues. The former measures the linear relation between the signal generated by a single tissue compartment and the acquired signal, while the latter measures the volume of single tissue compartment that is present in the voxel.

Given the known interdependence between the *T*_2_ times of tissues and the *T E* of the acquisition, some attempts at addressing this issue have been formulated making use of multi-TE multi-shell dMRI acquisitions (Veraart et al., 2018; Lampinen et al., 2020; Gong et al., 2020). Despite allowing to increase the signal-to-noise ratio (SNR) (Eichner et al., 2020), these techniques require a complete re-design of the experiments from acquisition to post-processing, posing severe limitations in terms of usability of already acquired data. This aspect is crucial in modern neuroimaging, where large studies like the Human Connectome Project (HCP) (Van Essen et al., 2012), the UK Biobank (Sudlow et al., 2015) and the Alzheimer Disease Neuroimaging Initiative (ADNI) (Mueller et al., 2005) invest significant amounts of time and financial resources to acquire data of large cohorts with standardised protocols that need to be carefully designed a priori.

In this work^1^ we will show that signal fractions are a biased estimation of volume fractions and that, under certain assumptions, the latter can still be retrieved from the first without acquiring new data or re-fitting the MC model. We call this new technique Multi-Tissue MC (MT-MC) model. To our knowledge MT-MC is the only general framework that allows to estimate *volume* fractions from single-TE multi-shell dMRI data. This novel formulation is inspired by the technique of Jeurissen et al. (2014) for the estimation of tissue-specific orientation distribution functions. The use of the MT-MC formulation solves some limitations of the previously mentioned multi-TE approaches and opens the door to the multi-tissue investigation of brain microstructure with data acquired with standard single-TE multi-shell dMRI protocols. Two algorithms for fitting the MT-MC model are proposed, one of which is designed to build on top of data already processed with standard MC models. Our new model is implemented and freely available in the Diffusion Microstructure Imaging in Python (Dmipy) (Fick et al., 2019) framework, which is an open source tool designed for the abstraction, simulation, and fitting of MC models of dMRI. The ability of the MT-MC model to retrieve the unbiased volume fractions is tested on both synthetic data generated with Dmipy and real data obtained from the HCP database.

This article is organized as follows: Section 2 is devoted to the theoretical aspects of MC modelling, highlighting why *signal* and *volume* fractions are not equivalent in general, and to the formalization of the proposed MT-MC model. In Section 3 we will present the design of the experiments and in Section 4 we will show the corresponding results, which are then discussed in Section 5, where also some conclusive remarks are presented.

## 2. Theory

### 2.1. Multi-Compartment models

Complex microstructural configurations can be modelled as a linear combination of few elementary compartments. For example, the diffusion within axons can be described as the motion of water molecules along a stick or within a cylinder, while diffusion in free water, like the one that can be observed in the CSF, can be modelled as an isotropic 3D Gaussian function. A vast portion of the dMRI literature of the last twenty years is devoted to the definition of compartmental models for the anisotropic intra-axonal and extra-axonal diffusivity and for the isotropic diffusivity. These are known as Multi Compartment (MC) models and they all describe the shape of the normalized dMRI signal *E* by means of the following linear combination of compartment-specific shapes:

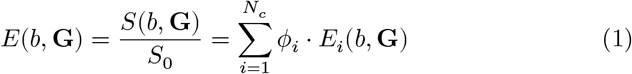

where *b* is the *b*-value, *S* is the raw diffusion signal, *S*_0_ is the diffusion signal acquired at *b* = 0, and **G** is the gradient direction, *N*_*c*_ is the number of considered compartments, *E*_*i*_ is the signal attenuation of compartment *i*, and *ϕ*_*i*_ is the portion of *E* explained by compartment *i*, i.e. the *signal fraction* of the compartment. The derivation of analytical expressions for the compartment-specific response functions has been researched broadly and deeply in the past literature. See the work of Panagiotaki et al. (2014) for a thorough review of the topic. Among the most used MC models we can mention the stick-and-ball model of Behrens et al. (2003), the ActiveAx model of Alexander et al. (2010) and the neurite orientation dispersion and density imaging (NODDI) model of Zhang et al. (2012). A generalized MC model has been proposed by Novikov et al. (2019) in what they called the *standard model* of dMRI in the brain.

The standard model is composed of three compartments which, borrowing the taxonomy from Panagiotaki et al. (2014), are defined as follows:

- The IC compartment is modelled as a stick whose free parameters are the parallel diffusivity *λ*_∥_ and the direction of the fiber population as the unit vector n. The corresponding signal is given by

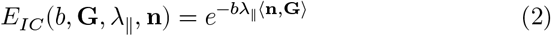

where ⟨**n, G**⟩ denotes the usual scalar product in ℝ^2^.
- The EC component is described by an axially symmetric Gaussian function (i.e., zeppelin), which can be defined as a diffusion tensor that depends on the parallel diffusivity *λ*_∥_, the perpendicular diffusivity *λ*_∥_ and the direction of the fiber population **n** (which is assumed to be the same as the one of the stick compartment). The signal shape is given by the classical tensor model

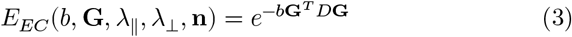

where the diffusion tensor is defined as *D* = (*λ*_∥_ − *λ*_⊥_) **nn**^*T*^ + *λ*_⊥_*I* and *I* is the 3-by-3 identity matrix.
- The CSF compartment is modelled an isotropic Gaussian function (i.e., ball), which is defined as a zeppelin with *λ*_∥_ = *λ*_⊥_= *λ*_*r*_ where *λ*_*r*_ is the radial diffusivity. The expression for the signal shape reads as follows:

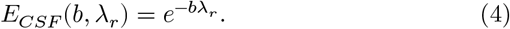 Notice that the first term in the definition of the diffusion tensor disappears, hence the model does not depend on the principal direction **n**, making the compartment *isotropic* as wanted.

Additionally, fiber dispersion is formalized as the convolution of the stick and zeppelin compartments with an ODF denoted by 𝒫. An example of such orientation function is the Watson distribution *W* (**n**, *κ*) (Mardia and Jupp, 1990), which assumes axial symmetry of the dispersion around the main direction of the bundle **n** ∈ 𝕊^2^ with concentration *κ*. The corresponding orientation dispersion index (ODI) can be computed as *ODI* = 2/*π* ⋅ arctan(1/*κ*) (Zhang et al., 2012).

Given the elements described in the previous lines, the MC formulation of the standard model is defined as

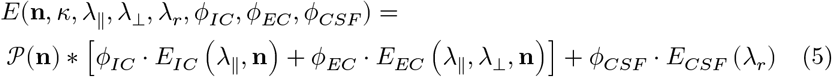

where * is spherical convolution operator and the dependence on the acquisition parameters *b* and **G** has been omitted for the sake of readability. Several constraints can be applied to the model given in Equation (5), among which the most commons are:

- the sum of the signal fractions is unitary: *ϕ*_*IC*_ + *ϕ*_*EC*_ + *ϕ*_*CSF*_ = 1;
- the perpendicular diffusivity of the EC compartment is tortuous (Szafer et al., 1995a,b), which in mathematical terms means

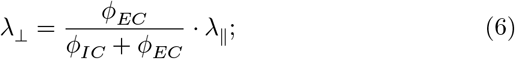

the parallel diffusivity of the IC and EC compartments is fixed (e.g. *λ*_∥_ = 1.7 ⋅ 10^−9^*m*^2^/*s* as in (Zhang et al., 2012));
- the radial diffusivity of the CSF compartment is fixed (e.g. *λ*_*r*_ = 3.0 ⋅ 10^−9^*m*^2^/*s* as in (Zhang et al., 2012)).

Recent studies questioned the validity of these constraints (Jelescu et al., 2016; Lampinen et al., 2017; Dell’Acqua and Tournier, 2019).

As highlighted by the left hand side of Equation (5), the model depends on eight parameters, where **n** is two-dimensional, yielding 9 degrees of freedom, to which one has to subtract the degrees of freedom covered by the constraints. The remaining parameters can be estimated solving the minimization problem

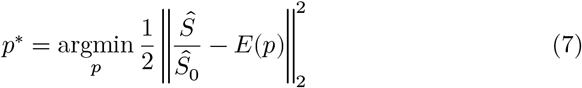

where *p* is the parameter vector, *Ŝ* is the acquired dMRI signal, *Ŝ*_0_ is the mean *b* = 0 image and *E*(*p*) is the realization of the forward model given in Equation (5). Fitting such parameters requires the acquisition of multi-shell data with at least one shell per compartment (Scherrer and Warfield, 2010). The obtained parameters *p*^*^ are the microstructural parameters that can finally be analysed for clinical or research purposes. In practice, the fitted signal fractions *ϕ*_*i*_ will likely not sum to 1, as they absorb any discrepancies between the normalised signal in the left hand side and the signal shapes in the left hand sides of Equation (5), in particular when more than one image is acquired at *b* = 0.

A thorough review on the variety of MC models of WM that can be defined with the current state-of-the-art tools is the one of Fick et al. (2019), where also the Dmipy package is presented. This software is the reference tool used throughout this work for the study of microstructure. More recently, MC models have been used to assess also the microstructural composition of the gray matter (GM) (Ganepola et al., 2018; Fukutomi et al., 2019; Villalon-reina et al., 2020), but the literature is still sparse and there is a lack of agreement on how to model the GM with MC models.

The key operation behind the definition of MC models is the division of the diffusion-weighted signal *S* by the non-diffusion-weighted component *S*_0_, which allows to retrieve the signal shape which is then modelled as the linear combination of the signal shape of the compartments that characterize the model. In the next section we are going to question the applicability of this division by *S*_0_.

### 2.2. MC models do not account for T2 differences

As stated in the previous section and formalised in Equation (1), MC models aim at fitting the signal shape *E* as the ratio of the PGSE signal *S* and the *S*_0_ amplitude. The implicit assumption that lies behind this formulation is that the *S*_0_ by which the acquired signal is divided is the same for all the modelled compartments. In particular, as the *S*_0_ image corresponds to the signal coming from the non-diffusion-weighted spin-echo sequence, we know that its amplitude depends on the echo time *T E* and the repetition time *T R* of the acquisition and on the *T*_1_ and *T*_2_ times of the sample. The relationship between these quantities reads as

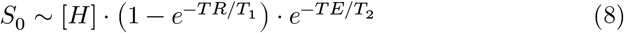

where [*H*] is the proton density in the sample. While in the formation of the *S*_0_ image the different *T*_1_ times of the tissues are negligible thanks to the length of *T R* (which is usually one order of magnitude longer than *T E*), tissues with different *T*_2_ will generate sensibly different contrast in the image (Plewes, 1994; Just and Thelen, 1988; Veraart et al., 2018). Figure 1 illustrates how this difference is visible in the *S*_0_ response of the WM and the CSF. These differences are the result of the different contrast in *T*_2_-weighted images between the different compartments. In order to understand how this difference in the *T*_2_ impacts the signal-fraction estimation, consider the following example. Let a voxel in the WM containing some partial volume of CSF, which is common in the corpus-callosum near the ventricles. In particular, let’s assume that the volume fractions are *f*_*WM*_ = 0.9 and *f*_*CSF*_ = 0.1. The corresponding signal equation will be

**Figure 1:**
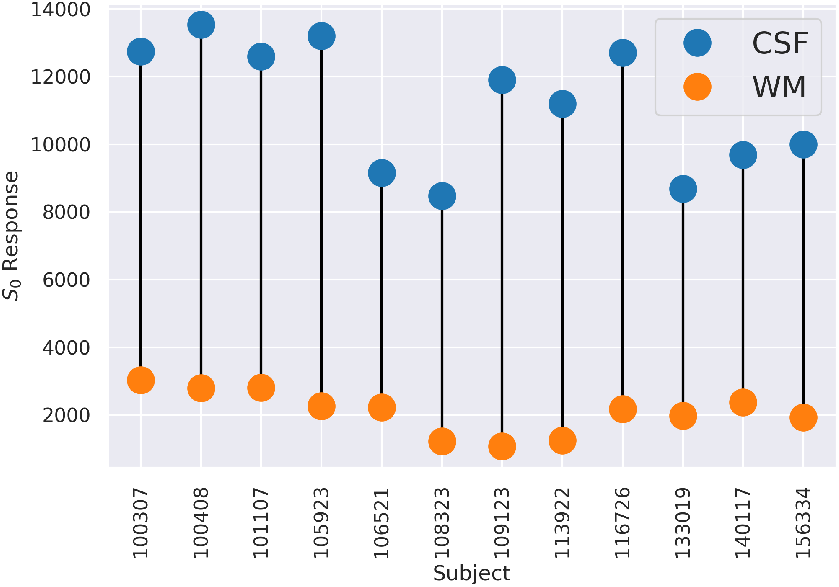
The figure shows the *S*_0_ response of the WM and of the CSF for twelve randomly picked subjects from the HCP database. Values are obtained with the heuristic technique of Dhollander et al. (2016) via Mrtrix3 (Tournier et al., 2019).

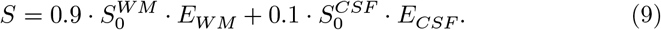

As highlighted by Figure 1, the value of 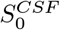 can be up to six times the one of 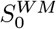. Including this into our toy model, hence defining 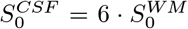, Equation (9) becomes

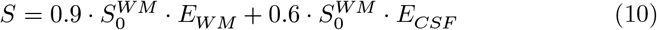

which after dividing both sides of the equation by the composite 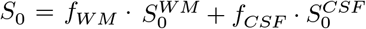 becomes

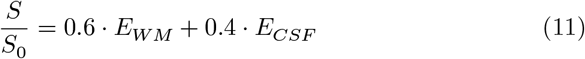

yielding the signal fractions *ϕ*_*WM*_ = 0.6 and *ϕ*_*CSF*_ = 0.4. This exampled showed how signal fractions and volume fractions are not interchangeable concepts when it comes to modelling multiple tissues having different *S*_0_ responses. Not taking into account this differences can lead to significant misrepresentations of the tissue composition, as showed in the previous example and in the results reported in Section 4.

### 2.3. Leveraging multi-TE sequences in Multi-Compartment modelling of the dMRI signal

If the problem of MC models is that they do not distinguish the *S*_0_ of different tissues because of the limitations of single-TE acquisition sequences like the one considered in the previous sections, the solution could simply be to use multi-TE (MTE) acquisitions, despite the required longer acquisition time. This idea has been investigated in recent works of Veraart et al. (2018), Lampinen et al. (2019, 2020), and Gong et al. (2020). These works are all based on the assumptions that the volume fraction of a tissue can not be computed with conventional multi-shell dMRI data acquired with a single echo time.

The TE-dependent Diffusion Imaging (TEdDI) technique proposed by Veraart et al. (2018) technique considers a rewriting of the MC equation that directly includes the contribution of the *T*_2_ time of the tissue modelled by the compartment and the *T E* of the acquisition into the volume fraction of each compartment. The same principles are followed in the more recent works of Lampinen et al. (2019, 2020) and of Gong et al. (2020). For the sake of coherence, we adapted the original notation used in the articles. The TEdDI model is designed to account for the *T E*/*T*_2_ effects in the same way as in the *S*_0_-image formation process described in Equation (8), obtaining

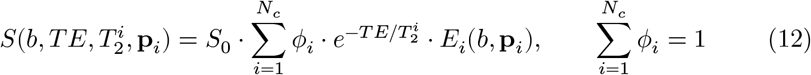

where 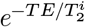 plays the role of the compartment-specific contribution of the *T*_2_ time and *S*_0_ is the proton density- and *T*_1_-weighted image, which corresponds to Equation (8) for *T E* = 0. Notice that the *T*_2_ time of each compartment is an independent variable of the model, hence it must be estimated in the fitting process. This requires the acquisition of multi-shell (to allow the use of multiple compartments) and multi-TE (to avoid degeneracy in the joint fitting of *ϕ*_*i*_ and 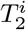) dMRI data. The *volume fraction* of each compartment is defined by Veraart et al. (2018) and Gong et al. (2020) as follows:

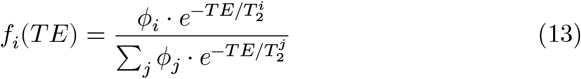

where one should notice how the volume fraction *f*_*i*_ depends on the echo time *T E*. Conversely, Lampinen et al. (2019, 2020) opted for defining the volume fractions as

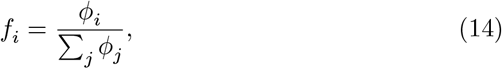

which corresponds to the normalisation of the *ϕ*_*i*_ retrieved from fitting the model given in Equation (12). The formulation provided in Equation (12) can be regarded as the multi-TE standard model of the dMRI signal in the human brain, in analogy with what reported by Novikov et al. (2019) (see Equation (5)).

Close variants of the MTE standard model have already been used in the previously cited works of Veraart et al. (2018), Lampinen et al. (2019, 2020) and Gong et al. (2020) to investigate the microstructure of the white matter of the brain. They showed that particular instances of the MTE standard model allow to assess how the *T*_2_ time of the acquired sample is formed by the different compartments. Also, with MTE-MC models they showed that the concept of *volume fraction* should not just be abandoned in favor of the concept of *signal fraction*. Its straightforward interpretability is of much appeal in brain pathology research (Suzuki et al., 2017; Hara et al., 2018; Vestergaard-Poulsen et al., 2007), where biomarkers are not only quantified but also contextualized, related to other non-microstructural information and interpreted.

Some limitations come with the use of such formulation. First, the volume fractions defined in Equation (13) are *T E*-dependent. This poses severe limitations in terms of usability and prevents from having a single index for the volume fraction of a compartment, which intuitively should be a characteristic of the sample, not of the acquisition. An additional limitation of MTE-MC modelling that we highlight is of methodological nature. Classical MC models are representations of the dMRI signal that rely on standard multi-shell acquisitions designed in a HARDI fashion which have been used in the last 15 years for the study of both microstructure and tractography-based structural connectivity. The MTE framework does have the merit to correct the signal/volume fraction ambiguity, but this is achieved by increasing the complexity of the acquisition, which requires multiple *T E* to be considered. For this reason, the MTE framework is not to be considered an alternative to the MC formulation but rather a new method for the estimation of microstructural parameters that spans the whole range from acquisition design to post-processing, preventing from correcting the estimation of volume fractions on datasets acquired in the past.

### 2.4. Multi-Tissue Multi-Compartment models

The standard formulation of MC models includes a normalization of the dMRI signal *S*_0_ by its non-diffusion-weighted component *S*_0_. This operation is performed in order to retrieve the shape *E* of the acquired signal. The shape is then modelled as a linear combination of signal shapes of different compartments. In Section 2.2 we showed how this formulation hides the assumption that all the tissues modelled by the compartments have the same *T*_2_ time (hence *S*_0_), highlighting how this is not true a-priori. The solutions to the multi-tissue problem proposed in the literature have the remarkable limitation of requiring the acquisition of multi-TE data to be used.

A solution to a similar problem has been proposed by Jeurissen et al. (2014) in the context of fODF estimation for multi-shell data, where the shell- and tissue-specific signal amplitude is leveraged in order to rescale the fODF that describes the signal shape of each considered tissue. This includes the response of each tissue in the *b* = 0 shell, hence the *S*_0_ of the tissues. The technique we are proposing builds on top of this idea. We highlight how similar solutions have been exploited also in the estimation of single-shell single-tissue response functions for the estimation of fODFs (Descoteaux et al., 2007; Tournier et al., 2007).

Let *N*_*c*_ be the number of compartments included in the model we want to design and let 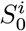 be the *S*_0_ response of compartment *i*. We define the *Multi-Tissue Multi-Compartment* (MT-MC) model as follows:

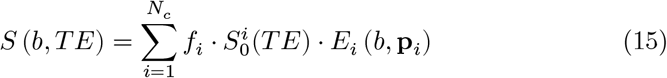

where *f*_*i*_ is the *volume fraction* of compartment *i* and 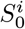 (*T E*) is the *S*_0_ response of the tissue modelled by compartment *i*. Notice that Equation (15) is equivalent to Equation (1) whenever 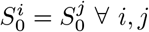, namely when all the tissues described by the MT-MC model have equal *S*_0_ responses.

In general, the signal fraction *ϕ*_*i*_ is not equivalent to the volume fraction *f*_*i*_ of the tissue modelled by the compartment. The only case in which they are equivalent is when all the tissues modelled by the MT-MC model have equal *S*_0_ responses. In that case, Equation (15) reduces to (1) after multiplying both the sides by *S*_0_. For this reason we say that *ϕ*_*i*_ is a *biased estimator* of *f*_*i*_. One could argue that the relationship between the signal fractions *ϕ*_*i*_ and the volume fractions *f*_*i*_ is just a rescaling, in which case the volume fractions could be retrieved with a simple correction that takes into account the *S*_0_ signal and the 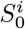 response of the compartment. This is always true, except when the volume fraction of the compartment is an independent variable in some other compartment. Using the tortuosity constraint to define the perpendicular diffusivity of the extra-axonal compartment, we establish a non-linear dependence between the compartmental fraction of the intra- and extra-cellular compartments. In this way, the diffusivity of the extra-axonal compartment, which has a non-linear relationship with the model, is defined as in Equation (6), forcing the intra- and extra-cellular signal/volume fractions to be non-linear parameters of the model. For instance, if the intra- and extra-axonal compartments have different *S*_0_, the perpendicular diffusivity of the EC computed with the tortuosity constraint defined on the signal fractions will be different from the one obtained from the volume fractions. As a consequence, two models defined with the two possible tortuosity constraints are not interchangeable and rescaling one’s volume fractions does not yield the other’s signal fractions.

#### 2.4.1. Fitting MT-MC models

The fitting of a MT-MC model is designed in a fashion similar to the one of MC models. Here we propose two different approaches. The first is a direct fitting that provides only the volume fractions (VF), while the second is a two-step strategy that builds on top of the fitting of the signal fractions and yields both the signal and the volume fractions (SVF), allowing to re-process in a MT fashion results that had previously been obtained on standard MC models. Given the acquired dMRI signal *S*, the corresponding *S*_0_, the number of compartments *N*_*c*_, the signal shape *E*_*i*_(**p**_*i*_) of compartment *i* depending on the parameter vector **p**_*i*_ and the compartment-specific signal amplitude 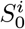, the fitting can be performed in the two following ways.

*VF*. The first approach directly fits the volume fractions by solving a least squares problem with respect to the microstructural parameters *f*_*i*_ and **p**_*i*_:

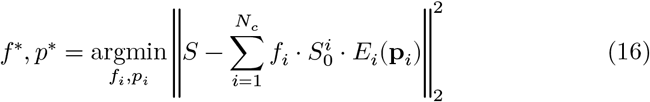

which can be solved with ordinary inverse-problem solvers. Here, the forward model is the one given in Equation (15). The procedure yields the volume fractions (VF) of the compartments.

*SVF*. The second approach extracts the volume fractions after fitting the signal fractions *ϕ*_*i*_ and the microstructural parameters **p**_*i*_ from the MC formulation of Equation (5). The volume fractions are retrieved as a rescaling of the signal fractions. The described procedure reads as follows:

1. Solve the associated MC problem:

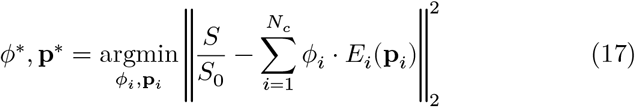

where the product of the minimization problem is the signal fraction *ϕ*_*i*_and the parameter vector **p**_*i*_ of each compartment *i*;
2. Fix the fitted non-signal-fraction parameters in the MT-MC model. At this point the volume fractions are not related to each other (or to other compartments in general) and it is therefore possible to estimate them by rescaling the signal fractions. The rescaling is the one suggested by the comparison of the coefficients that multiply the signal shapes in Equations (1) and (15) and reads as follows:

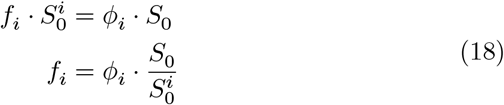

yielding a simple operation that allows to retrieve volume fractions from signal fractions once the *S*_0_ of each compartment is known. Both the signal and volume fractions of each compartment are returned.

To employ either of the two fitting strategies, extra caution must be taken towards the use of the tortuosity constraint. The intra- and extra-axonal fractions used for the definition of the perpendicular diffusivity can be either the signal fractions or the volume fractions of the compartments, i.e.,

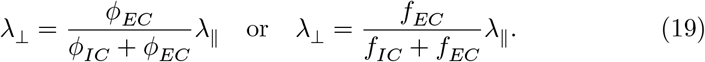

The choice influences the whole model design and can not be reverted in the fitting process. In particular, switching between signal fractions and volume fractions with the 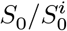 rescaling must be done keeping in mind that the tortuous parameters have been obtained using a specific type of fraction, and the results should be interpreted accordingly. In an effort to keep the notation coherent with the previous literature, we will say that whenever the tortuosity constraint is defined using the volume fractions *f*_*i*_ we will have a *MT-corrected* tortuosity constraint.

The SVF strategy is the one implemented in Dmipy (Fick et al., 2019), which to our knowledge is the only available framework for generalised MC modelling that includes the definition of MT-MC models. As far as specific instances of MT-MC models are concerned, the multi-shell multi-tissue CSD technique of (Jeurissen et al., 2014) is implemented in Mrtrix3 (Tournier et al., 2019).

### 2.5. The MT Standard Model of dMRI in White Matter

In this Section, we define a MT generalization of the standard model of dMRI in WM as described by Novikov et al. (2019). We recall that the model includes a stick and a zeppelin compartment for the intra- and extra-cellular diffusivity respectively and a ball that accounts for the isotropic diffusivity in the CSF and other isotropic structures. Let 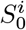 be the *S*_0_ response of the tissue modelled by each compartment *i* and 𝒫 : 𝕊^2^ → ℝ^+^ the orientation distribution. The MT standard model of dMRI in WM is given by

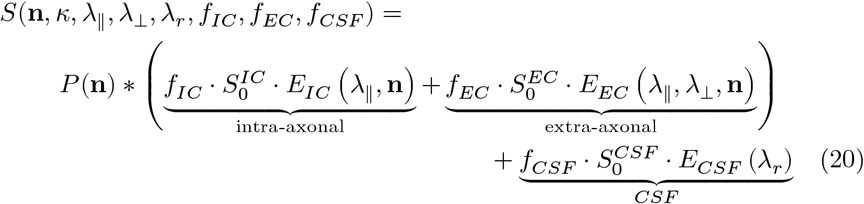

where the compartment specific parameters are defined as in Section 2.1. Three scenarios can be described with this model:

- *3-tissue model* - The three compartments describe tissues with distinct *S*_0_ responses. This corresponds to the explicit case of Equation (20).
- *2-tissue model* - The two anisotropic compartments model tissues whose *S*_0_ is equal. Typically, it is the *S*_0_ of the WM, so we say that 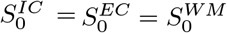 and 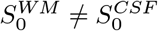.
- *1-tissue model* - In absence of any prior knowledge on the *S*_0_ of the three tissues, they are considered all equal. We denote this as 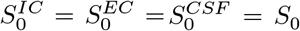 where *S*_0_ is the average across the images acquired at *b* = 0 *s*/*mm*^2^.

Notice that the 1-tissue scenario is mathematically equivalent to the single-tissue (ST) standard model of Equation (20), hence in that case the volume fractions are equivalent to the signal fractions.

## 3. Methods

### 3.1. Dataset

#### 3.1.1. Synthetic data

The simulated dataset is obtained from the forward model given by Equation (20) and generated with Dmipy (Fick et al., 2019). A total of 10000 voxels was simulated on a multi-shell acquisition scheme identical to the one that will be considered on the real dataset, which includes a TE of 0.0895*s* and is composed of 288 samples subdivided in 18 points at *b* = 0*s*/*mm*^2^ and 90 diffusion-weighted samples obtained with uniformly distributed directions at *b* = 1000*s*/*mm*^2^, *b* = 2000*s*/*mm*^2^ and *b* = 3000*s*/*mm*^2^ for a total of 3 diffusion-weighted shells plus the *b* = 0 shell. The direction of the two anisotropic compartments was set to n = [0, 0] ∈ 𝕊^2^ for all the voxels. The *T*_2_ time of each tissue was randomly sampled from a uniform distribution in the range specified in Table 1. The corresponding *S*_0_ was then computed as 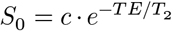 where *c* is a scaling parameter that positions the value of *S*_0_ in a realistic range and we tuned to *c* = 1400. The ODI of the Watson distribution was sampled from a uniform distribution in the range specified in Table 1. Finally, the volume fractions of each compartment were randomly generated from a uniform distribution in the range specified in Table 1, then normalized in such a way that their sum was equal to 1. The choice of each range was tuned to mimic the single-bundle configuration in the WM that one expects to be able to model with the considered formulation. An additive rician noise was added to the simulated data to obtain a signal-to-noise ratio equal to 30.

**Table 1:** For each parameter used in the definition of the forward model of the synthetic dataset we report the minimum and maximum value of the uniform distribution from which it was drawn.

| Parameter | Min | Max |
| --- | --- | --- |
| $f_{IC}$ | 0.5 | 0.8 |
| $f_{EC}$ | 0.3 | 0.5 |
| $f_{CSF}$ | 0.3 | 0.7 |
| $T_2^{IC}$ | 0.080s | 0.100s |
| $T_2^{EC}$ | 0.050s | 0.070s |
| $T_2^{CSF}$ | 0.900s | 1.100s |
| $ODI$ | 0.02 | 0.99 |

#### 3.1.2. Real data

From the Human Connectome Project (HCP) database we considered three randomly picked subjects^2^ available at the Connectome Coordination Facility (Van Essen et al., 2012; Sotiropoulos et al., 2013). For each subject a total of 288 images is acquired, subdivided in 18 volumes at *b* = 0*s*/*mm*^2^ and 90 diffusion-weighted volumes obtained at uniformly distributed directions at *b* = 1000*s*/*mm*^2^, *b* = 2000*s*/*mm*^2^ and *b* = 3000*s*/*mm*^2^ for a total of 3 shells. All subjects provided written informed consent, procedures were approved by the ethics committee and the research was performed in compliance with the Code of Ethics of the World Medical Association (Declaration of Helsinki).

To our knowledge, current state-of-the-art techniques do not allow to estimate subject-specific *S*_0_ responses of the IC and EC compartments, while the *S*_0_ of the CSF compartment can be estimated together with 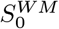 via techniques such as the heuristic approach of Dhollander et al. (2016). For this reason, we analysed the aforementioned data with a 1-tissue and a 2-tissue model where 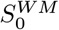 and 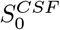 have been estimated with the Dhollander technique. The obtained values of *S*_0_ are displayed in Figure 1 and reported in Table 2.

**Table 2:** *S*_0_ response of the WM and of the CSF of the three studied HCP subjects. Values are obtained with the heuristic technique of Dhollander et al. (2016) via Mrtrix3 (Tournier et al., 2019).

| Subject | $S_0^{WM}$ | $S_0^{CSF}$ |
| --- | --- | --- |
| #1 | 3024 | 12740 |
| #2 | 2794 | 13531 |
| #3 | 2811 | 12598 |

### 3.2. Model fitting

The model considered in the performed experiments is the MT standard model defined in the previous section with fixed 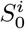 and three additional constraints:

- The fiber orientation distribution is modelled with a Watson distribution of axis **n** and fixed ODI.
- The perpendicular diffusivity is subject to the tortuosity constraint, hence

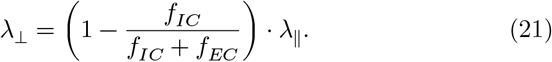
- The parallel and radial diffusivity are fixed to *λ*_∥_ = 1.7 ⋅ 10^−9^*m*^2^*s*^−1^ and *λ*_*r*_ = 3.0 ⋅ 10^−9^*m*^2^*s*^−1^ respectively.

The free parameters that are left are [*f*_*IC*_, *f*_*EC*_, *f*_*ISO*_, **n**, *κ*], where we recall that the unit vector **n** is expressed in spherical coordinates [*θ, φ*]. the fitting was performed with Dmipy (Fick et al., 2019, version 1.0.3) using the SVF procedure described in Section 2.4.1 in order to retrieve both the signal fractions and the volume fractions to be compared.

The difference between the absolute errors obtained with each instance of the model is tested with a Wilcoxon signed-rank test (Wilcoxon, 1945) with *α* = 0.05 and measured with the rank biserial correlation, i.e., an effect size measure in the [0, 1] range.

## 4. Results

### 4.1. Synthetic data

We fitted the volume fractions 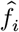 of each compartment with the SVF procedure explained in Section 2.4.1 with the 1-tissue, 2-tissue and 3-tissue model, both with standard and MT-corrected tortuosity. The latter is actually different from the standard tortuosity only when the 3-tissue model is considered. When the 2-tissue model is employed, the 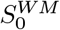 response is computed as the weighted average of the *S*_0_ response of the IC and EC compartments, hence

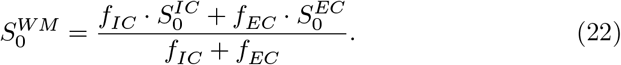

Once each 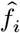 was estimated, we computed the absolute fitting error 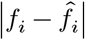. The difference between the absolute errors obtained with each model is tested with a Wilcoxon signed-rank test (Wilcoxon, 1945) with *α* = 0.05 and measured with the rank biserial correlation, i.e., an effect size measure in the [0, 1] range. In Figure 2 we report the boxplot of the distribution of the absolute fitting error across 10000 simulations (top row) and the rank biserial correlation (bottom row). As we expected, the signal fractions retrieved by the 1-tissue model are biased estimates of the volume fractions retrieved with the 2-tissue and 3-tissue model. The bias in the estimation of the volume fraction of the CSF compartment is four times bigger than the one of the IC and EC compartment. This is coherent with the fact that the *S*_0_ of the CSF compartment is much higher than the one of the IC and EC compartments. The error decreases importantly when the 2-tissue model is used. Here, the IC volume fraction has absolute error comparable to the one of the 3-tissue models. The first factor that could induce such phenomenon is the definition of 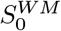, which by design of the experiment will be closer to the *S*_0_ of the IC than to the one of the EC compartment (*f*_*IC*_ > *f*_*EC*_ as reported in Table 1). This induces the estimated EC volume fraction to be farther from the ground truth than the one of the IC compartment. This difference is reflected in the absolute error of the estimated volume fraction of the CSF compartment, which is affected by the presence of the non-zero perpendicular diffusivity of the EC compartment. Nevertheless, the estimation error of the CSF volume fraction is much lower than in the 1-tissue model thanks to the inclusion of the specific 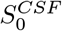 in the formulation. Finally, the 3-tissue model retrieves volume fractions that are in line with the ground truth ones. A notable aspect concerns the inclusion of the MTT correction, which is shown to significantly improve the estimation of the IC volume fraction. This improvement corresponds to a deterioration (albite lower in scale) of the estimation of the EC volume fraction. The estimation of the CSF volume fraction does not change significantly when the MTT correction is employed.

**Figure 2:**
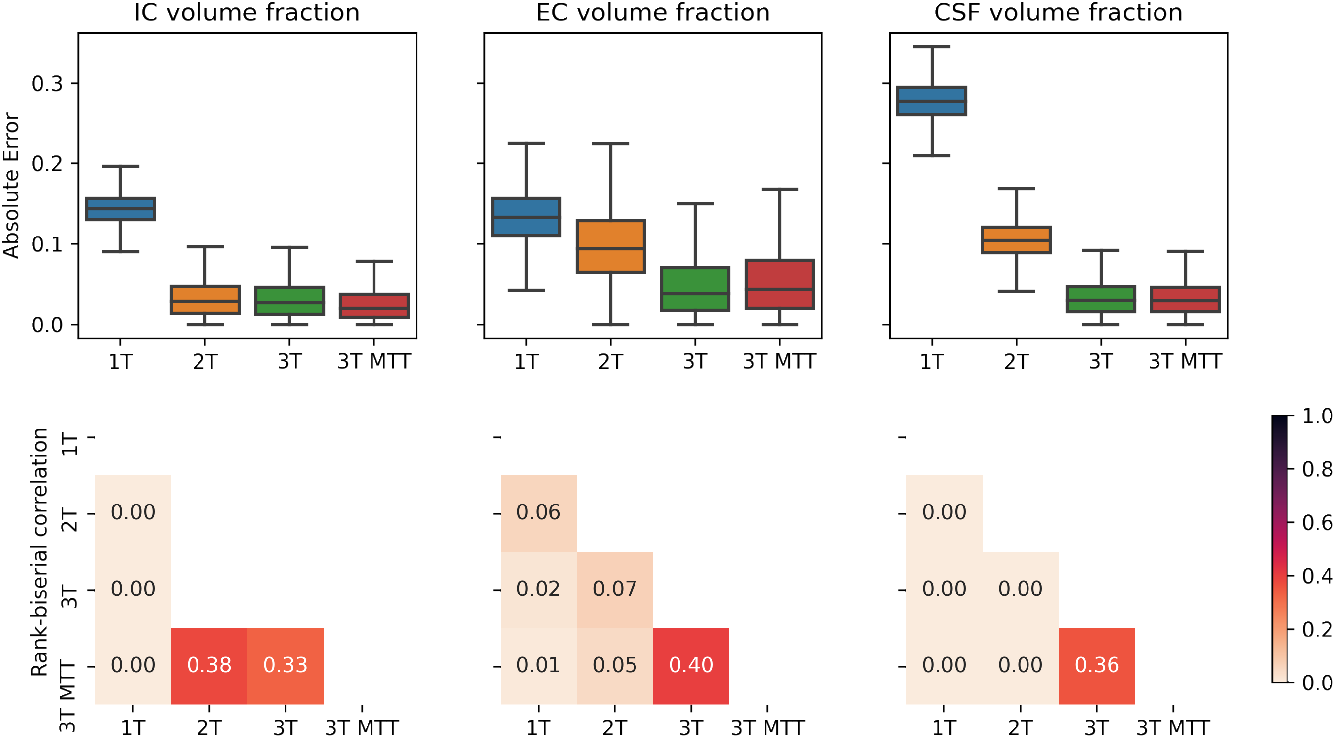
The first row shows the boxplot of the absolute error of the estimated volume fraction of each compartment computed on the synthetic dataset. The 1T categorical variable corresponds to the 1-tissue model, 2T to the 2-tissue, 3T to the 3-tissue with standard tortuosity and 3T MTT to the 3-tissue model with MT-corrected Tortuosity (MTT). The second row shows, for each compartment, the rank biserial correlations measuring the difference between the absolute errors of each model obtained from a Wilcoxon test with *α* = 0.05: the lower the correlation value, the higher the difference between the underlying distributions. Only statistically significant results are reported.

### 4.2. Real data

For each model, we fitted the signal and volume fractions with the SVF technique. Figure 3 shows the distribution of the signal fraction and volume fraction of each compartment in the WM for three HCP subjects. The WM mask was computed with FSL *fast* from the *T*_1_-weighted image with 1.25*mm* voxel size available at the HCP database, then dilated by one voxel with Mr-trix3’s (Tournier et al., 2019) *maskfilter* command to smooth the boundary. For each subject, the difference between the distribution of the signal and the volume fractions is tested with a Wilcoxon signed-rank test (Wilcoxon, 1945) with *α* = 0.05 and measured with the rank biserial correlation, i.e., an effect size measure in the [0, 1] range. The results of this test are reported in Table 3.

**Table 3:** For each subject (1, 2, 3) and compartment (IC, EC, CSF), the table displays the value of the rank biserial correlation *r* and the corresponding p-value *p* computed obtained from a Wilcoxon signed rank test (Wilcoxon, 1945) with *α* = 0.05. The showed values are computed with Scipy (Harris et al., 2020) and all the performed comparisons exhibit statistically significant differences.

| Subject | IC |  | EC |  | CSF |  |
| --- | --- | --- | --- | --- | --- | --- |
| | $r$ | $p$ | $r$ | $p$ | $r$ | $p$ |
| #1 | 0.243 | 0.0 | 0.036 | 0.0 | 0.002 | 0.0 |
| #2 | 0.221 | 0.0 | 0.142 | 0.0 | 0.002 | 0.0 |
| #3 | 0.197 | 0.0 | 0.100 | 0.0 | 0.001 | 0.0 |

**Figure 3:**
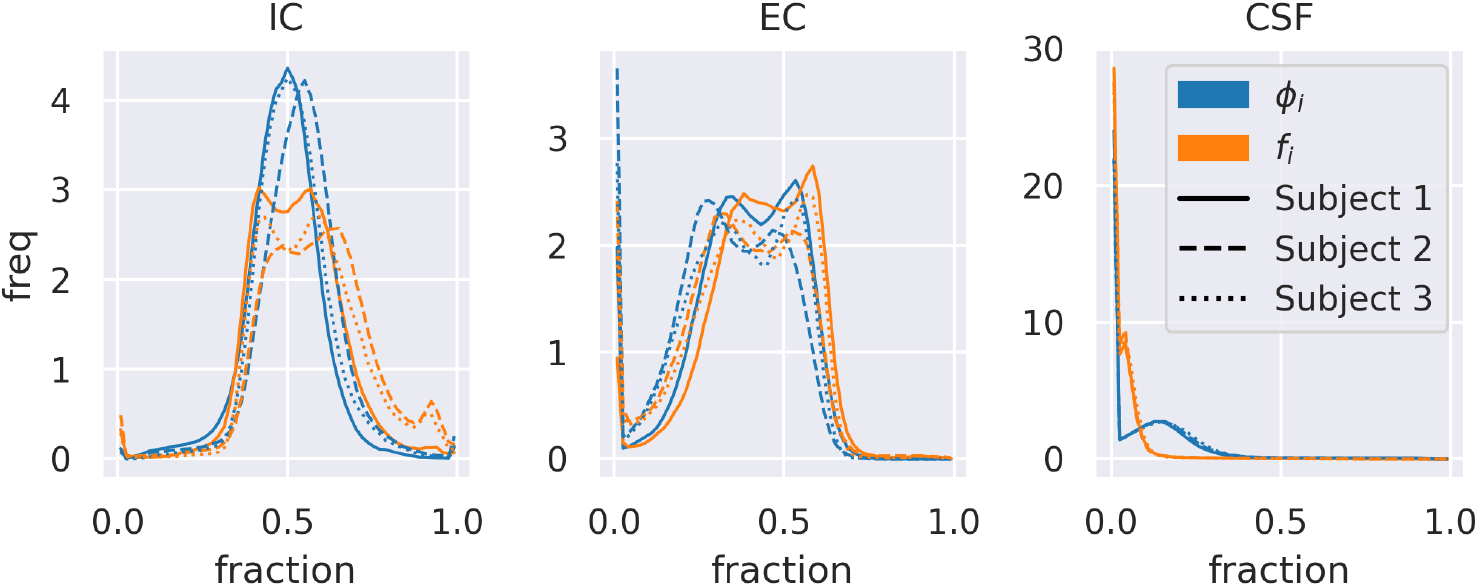
The displayed data are obtained three subjects of the HCP database (solid lines, dashed lines, and dotted lines). Each panel shows the distribution of the signal fraction and the volume fraction of the IC, EC and CSF compartments respectively. The blue lines correspond to *signal* fractions and the orange lines to *volume* fractions.

We recall that we considered a 2-tissue model by compressing the IC and EC compartments in a unique block that describes the WM tissue. The distribution of the volume fractions of the IC and EC compartments showed in Figure 3 is right-shifted with respect to the distribution of the corresponding signal fractions. On the contrary, the distribution of CSF volume fractions in the WM mask is shifted towards lower values with respect to the corresponding signal fractions. This means that the signal fraction *underestimates* the presence of the intracellular compartment in favour of the CSF compartment. This behaviour is consisted in all the tested subjects.

This is coherent with the proportion between 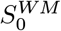 and 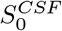, as the former is typically lower than the acquired *S*_0_ and the latter is higher. The results displayed in Figure 4 show how, within the WM mask, the WM volume fraction is globally higher than the WM signal fraction. Also, the absolute difference between the two exhibits some uniformity within the considered sample. The macroscopic differences between the left and right hemispheres present in all the three subjects may be due to some bias field effect that we did not include in the model and survived the minimal preprocessing of the data (Glasser et al., 2013).

**Figure 4:**
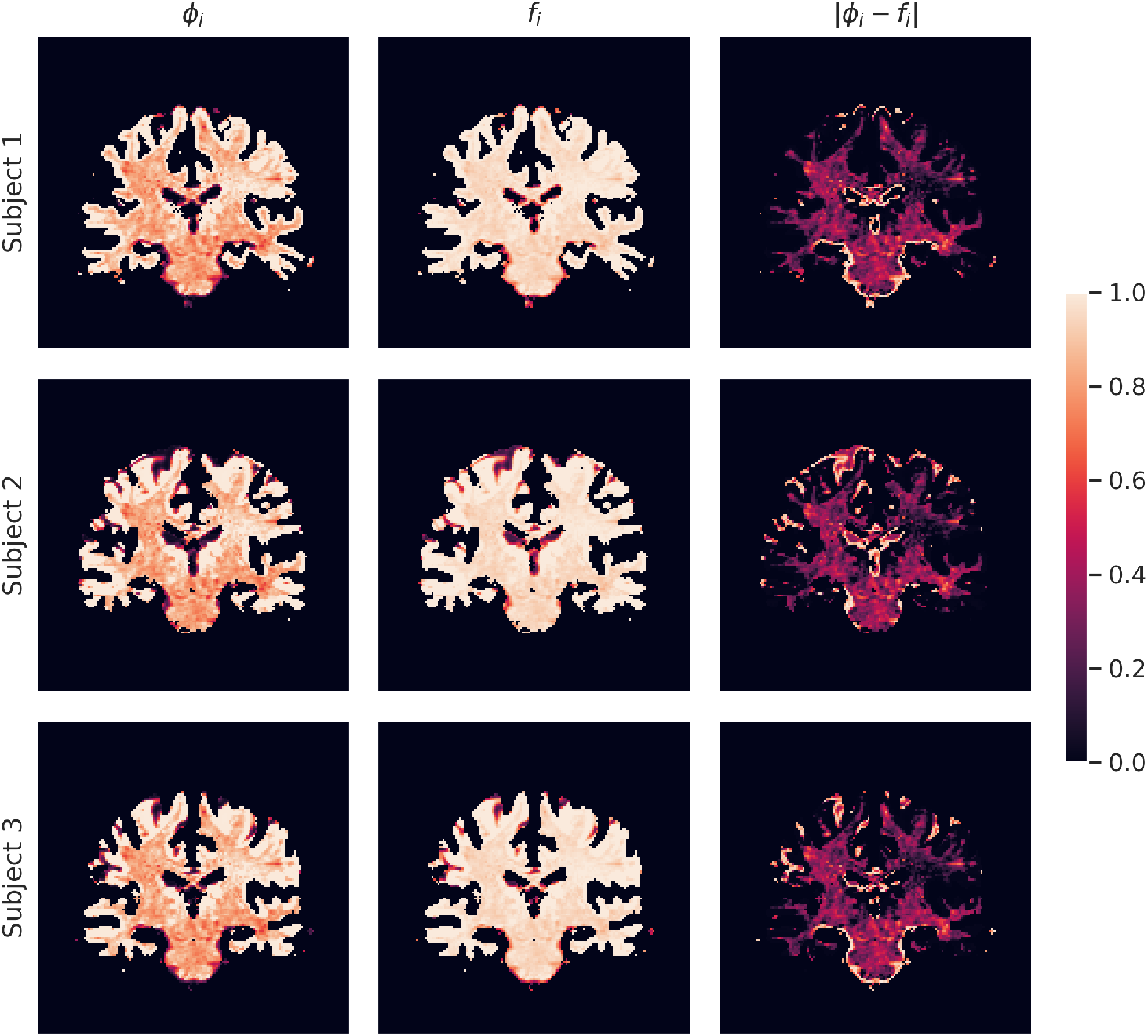
Signal fractions (first column), volume fractions (second column), and their absolute difference (third column) for three subjects of the HCP database. Brighter colors correspond to higher fractions (in the first two columns) and errors (in the third column). Voxels shown in orange/red and black correspond to decreasingly lower values of the same fractions and errors.

## 5. Discussion

In this paper we analysed how multi-compartment models of brain tissue microstructure can be adapted to account for the presence of tissues having different *T*_2_ relaxation times. In particular, we focused on the capability of such models to estimate the volume fraction of each tissue in the WM. We proposed a solution based on single-TE dMRI data, in contrast with the state-of-the-art techniques that require multi-TE dMRI data. Our results on both synthetic and in-vivo data show that *signal fraction* and *volume fraction* are not interchangeable concepts in the context of MC microstructure modelling. The shift of paradigm from signal fractions to volume fractions has already been shown to improve the estimation of fODFs (Jeurissen et al., 2014) and in this work we transferred the same approach to the field of MC models of brain tissue microstructure, leveraging the differences between the *S*_0_ responses of each modelled tissue.

Overall, the presented results yielded an empirical confirm of the theoretical considerations made in this work. In particular, the following aspects are highlighted:

- With *single-TE dMRI data* it is possible to retrieve tissue-specific volume fractions. Under the assumption that the IC and EC compartments have equal *S*_0_ response, techniques like the one of Dhollander et al. (2016) allow to define the 2-tissue model used in this section, opening the door to a better estimation of the compartment-specific volume fractions. This is made possible by the MT-version of the standard model of dMRI in the WM that we presented in this work. It models multiple tissues in a MC fashion *without* requiring multi-TE acquisition, which are conversely necessary in order to use other state-of-the-art models.
- *Signal fractions and volume fractions are not equivalent in general*. This fact has considerable implications in clinical context. Previous studies that drew conclusions based on the idea of inspecting volume fractions with single-TE dMRI need to be re-interpreted in light of the fact that what they are based on is the *signal* fraction of the tissues and not their volume fraction. How those differences are expressed in the presence of pathology or group differences remains unexplored and needs to be assessed in future studies.

We designed a multi-tissue version of the standard model of dMRI in the WM, which allows to separate the contribution of the intra-axonal, the extra-axonal and the CSF compartments and estimate the corresponding three volume fractions. The results reported in Figure 2 suggest that 2-tissue and 3-tissue models are always preferable to the 1-tissue model. A bigger improvement is obtained by considering two tissues instead of one, compared to the shift from the 2-tissue to the 3-tissue model. This is due to the proportion between the *S*_0_s of each tissue, which sees 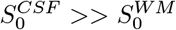, with 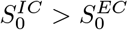 but the latter difference is lower than the former (Jeurissen et al., 2014).

A remarkable property of the proposed MT-MC model is that not only it can be straightforwardly fitted on single-TE dMRI data (VF strategy), but it can also re-use the results obtained with the MC version of the model (which in principle would have returned only the signal fraction of each compartment) and yield the volume fractions by means of an elementary rescaling operation (SVF strategy). While employing the SVF solution, extra care must be devoted to the use of the tortuosity constraint. Rescaling signal fractions obtained using the non-MT-corrected tortuosity constraint yields the volume fractions of a model where the perpendicular diffusivity of the EC compartment has been obtained using signal fractions, configuring an ambiguous (if not degenerate) solution.

The proposed model strongly relies on the external estimation of the *T*_2_ or the *S*_0_ of the modelled tissues. Our experiments on real data leveraged the heuristic of Dhollander et al. (2016) to retrieve the *S*_0_ of the WM and the CSF. Understanding how this choice affects the estimation of volume fractions is out of the scope of this work, but the raised question suggests that further efforts should be devoted to researching techniques that estimate tissue-specific *S*_0_ responses using single-TE data. Additionally, analysing the proportion between the *S*_0_ of each tissue in a large cohort of subjects could highlight patterns that could be exploited. If hypothetically the *T*_2_ relaxation time of extra-axonal compartment was showed to be a constant fraction of the *T*_2_ of the intra-axonal compartment, this could straightforwardly be encoded in the model.

The difference between signal fractions and volume fractions has implications also in the field of tractography filtering (Frigo et al., 2020a), where a coefficient is assigned to each streamline in a tractogram weighing its contribution to the formation of the dMRI signal. In the COMMIT framework (Daducci et al., 2015) these coefficients are the signal fractions associated to each streamline. The model can be easily adapted to obtain the volume fraction associated to each streamline, in particular in the context of the recent work of Barakovic et al. (2020), where streamlines are associated to bundle-specific *T*_2_ times.

## 6. Conclusion

In this work, we analyzed the brain tissue microstructure estimation via multi-compartment models of dMRI. We tackled the known limitation concerning the inability of state-of-the-art multi-compartment models to describe multiple tissues having distinct *T*_2_ relaxation times. We showed how what has always been considered the volume fraction of a certain tissue is actually the signal fraction of the same tissue. State-of-the-art techniques for overtaking such limitation rely on multi-TE dMRI data. Here, we introduced the Multi-Tissue Multi-Compartment models of dMRI, which allow to model multiple tissues at the same time using single-TE dMRI data. Moreover, we formulated a generalised multi-tissue modelling framework that encompasses both single-TE and multi-TE multi-tissue models. Our results indicate that with single-TE dMRI data alone one can model multiple tissues at the same time using the proposed multi-tissue multi-compartment models.

## Acknowledgements

This work was funded by the European Research Council (ERC) under the European Union’s Horizon 2020 research and innovation program (ERC Advanced Grant agreement No 694665: CoBCoM - Computational Brain Connectivity Mapping). Data were provided by the Human Connectome Project, WU-Minn Consortium (Principal Investigators: David Van Essen and Kamil Ugurbil; 1U54MH091657) funded by the 16 NIH Institutes and Centers that support the NIH Blueprint for Neuroscience Research; and by the McDonnell Center for Systems Neuroscience at Washington University. This work has been supported by the French government, through the 3IA Côte D’Azur Investments in the Future project managed by the National Research Agency (ANR) with the reference number ANR-19-P3IA-0002.

## Declaration of competing interest

The authors declare that they have no known competing financial interests or personal relationships that could have appeared to influence the work reported in this paper.

## Open Science

In this work we used only publicly available data from the Human Connectome Project database (Van Essen et al., 2012) and open source code from the Dmipy (Fick et al., 2019, version 1.0.3) Python package and the Mrtrix3 (Tournier et al., 2019) suite.

## Footnotes

1 This work has partially been presented at the International Symposium on Biomedical Imaging of 2020 (Frigo et al., 2020b) and at the 26th meeting of the Organization for Human Brain Mapping (Frigo et al., 2020c).

2 ID subject 1: 100307, ID subject 2: 100408, ID subject 3: 101107.

## Notes

### Competing Interest Statement

The authors have declared no competing interest.

### Summary of Updates

General rephrasing + missing references and typos fixing; improved presentation of the results; clarification of the theory

## References

Afzali, M., Pieciak, T., Newman, S., Garifallidis, E., Özarslan, E., Cheng, H., Jones, D.K., 2020. The sensitivity of diffusion mri to microstructural properties and experimental factors. Journal of Neuroscience Methods, 108951.

Alexander, D.C., Hubbard, P.L., Hall, M.G., Moore, E.A., Ptito, M., Parker, G.J., Dyrby, T.B., 2010. Orientationally invariant indices of axon diameter and density from diffusion mri. Neuroimage 52, 1374–1389.

Assaf, Y., Basser, P.J., 2005. Composite hindered and restricted model of diffusion (charmed) mr imaging of the human brain. Neuroimage 27, 48–58.

Assaf, Y., Blumenfeld-Katzir, T., Yovel, Y., Basser, P.J., 2008. Axcaliber: a method for measuring axon diameter distribution from diffusion mri. Magnetic Resonance in Medicine: An Official Journal of the International Society for Magnetic Resonance in Medicine 59, 1347–1354.

Barakovic, M., Tax, C.M., Rudrapatna, U.S., Chamberland, M., Rafael-Patino, J., Granziera, C., Thiran, J.P., Daducci, A., Canales-Rodríguez, E.J., Jones, D.K., 2020. Resolving bundle-specific intra-axonal t2 values within a voxel using diffusion-relaxation tract-based estimation. NeuroImage, 117617.

Behrens, T.E., Woolrich, M.W., Jenkinson, M., Johansen-Berg, H., Nunes, R.G., et al., 2003. Characterization and propagation of uncertainty in diffusion-weighted mr imaging. Magnetic Resonance in Medicine: An Official Journal of the International Society for Magnetic Resonance in Medicine 50, 1077–1088.

Daducci, A., Dal Palù, A., Lemkaddem, A., Thiran, J.P., 2015. Commit: convex optimization modeling for microstructure informed tractography. IEEE transactions on medical imaging 34, 246–257.

Dell’Acqua, F., Tournier, J.D., 2019. Modelling white matter with spherical deconvolution: How and why? NMR in Biomedicine 32, e3945.

Descoteaux, M., Angelino, E., Fitzgibbons, S., Deriche, R., 2007. Regularized, fast, and robust analytical q-ball imaging. Magnetic Resonance in Medicine 58, 497–510. URL: https://doi.org/10.1002/mrm.21277, doi:10.1002/mrm.21277.

Dhollander, T., Raffelt, D., Connelly, A., 2016. Unsupervised 3-tissue response function estimation from single-shell or multi-shell diffusion mr data without a co-registered t1 image, in: ISMRM Workshop on Breaking the Barriers of Diffusion MRI, p. 2016.

Eichner, C., Paquette, M., Mildner, T., Schlumm, T., Pléh, K., Samuni, L., Crockford, C., Wittig, R.M., Jäger, C., Möller, H.E., et al., 2020. Increased sensitivity and signal-to-noise ratio in diffusion-weighted mri using multi-echo acquisitions. Neuroimage.

Farooq, H., Xu, J., Nam, J.W., Keefe, D.F., Yacoub, E., Georgiou, T., Lenglet, C., 2016. Microstructure imaging of crossing (mix) white matter fibers from diffusion mri. Scientific reports 6, 1–9.

Fick, R., Wassermann, D., Deriche, R., 2019. The dmipy toolbox: Diffusion mri multi-compartment modeling and microstructure recovery made easy. Frontiers in Neuroinformatics 13. doi:/10.3389/fninf2019.00064.

Frigo, M., Deslauriers-Gauthier, S., Parker, D., Ismail, A.A.O., Kim, J.J., Verma, R., Deriche, R., 2020a. Diffusion mri tractography filtering techniques change the topology of structural connectomes. Journal of Neural Engineering 17, 065002.

Frigo, M., Fick, R., Zucchelli, M., Deslauriers-Gauthier, S., Deriche, R., 2020b. Multi tissue modelling of diffusion mri signal reveals volume fraction bias, in: 2020 IEEE 17th International Symposium on Biomedical Imaging (ISBI), IEEE. pp. 991–994.

Frigo, M., Zucchelli, M., Fick, R., Deslauriers-Gauthier, S., Deriche, R., 2020c. Multi-compartment modelling of diffusion mri signal shows te-based volume fraction bias, in: OHBM 2020-26th meeting of the Organization of Human Brain Mapping, pp. hal–02925963.

Fukutomi, H., Glasser, M.F., Murata, K., Akasaka, T., Fujimoto, K., Yamamoto, T., Autio, J.A., Okada, T., Togashi, K., Zhang, H., et al., 2019. Diffusion tensor model links to neurite orientation dispersion and density imaging at high b-value in cerebral cortical gray matter. Scientific reports 9, 1–12.

Ganepola, T., Nagy, Z., Ghosh, A., Papadopoulo, T., Alexander, D.C., Sereno, M.I., 2018. Using diffusion mri to discriminate areas of cortical grey matter. NeuroImage 182, 456–468.

Glasser, M.F., Sotiropoulos, S.N., Wilson, J.A., Coalson, T.S., Fischl, B., et al., 2013. The minimal preprocessing pipelines for the human connectome project. Neuroimage 80, 105–124.

Gong, T., Tong, Q., He, H., Sun, Y., Zhong, J., Zhang, H., 2020. Mtenoddi: Multi-te noddi for disentangling non-t2-weighted signal fractions from compartment-specific t2 relaxation times. NeuroImage, 116906.

Hara, S., Hori, M., Murata, S., Ueda, R., Tanaka, Y., Inaji, M., Maehara, T., Aoki, S., Nariai, T., 2018. Microstructural damage in normal-appearing brain parenchyma and neurocognitive dysfunction in adult moyamoya disease. Stroke 49, 2504–2507.

Harris, C.R., Millman, K.J., van der Walt, S.J., Gommers, R., Virtanen, P., Cournapeau, D., Wieser, E., Taylor, J., Berg, S., Smith, N.J., Kern, R., Picus, M., Hoyer, S., van Kerkwijk, M.H., Brett, M., Haldane, A., Fernández del Río, J., Wiebe, M., Peterson, P., Gérard-Marchant, P., Sheppard, K., Reddy, T., Weckesser, W., Abbasi, H., Gohlke, C., Oliphant, T.E., 2020. Array programming with NumPy. Nature 585, 357–362. doi:10.1038/s41586-020-2649-2.

Jelescu, I.O., Budde, M.D., 2017. Design and validation of diffusion mri models of white matter. Frontiers in physics 5, 61.

Jelescu, I.O., Veraart, J., Fieremans, E., Novikov, D.S., 2016. Degeneracy in model parameter estimation for multi-compartmental diffusion in neuronal tissue. NMR in Biomedicine 29, 33–47.

Jeurissen, B., Tournier, J.D., Dhollander, T., Connelly, A., Sijbers, J., 2014. Multi-tissue constrained spherical deconvolution for improved analysis of multi-shell diffusion mri data. NeuroImage 103, 411–426.

Just, M., Thelen, M., 1988. Tissue characterization with t1, t2, and proton density values: results in 160 patients with brain tumors. Radiology 169, 779–785.

Kaden, E., Kelm, N.D., Carson, R.P., Does, M.D., Alexander, D.C., 2016. Multi-compartment microscopic diffusion imaging. NeuroImage 139, 346–359.

Lampinen, B., Szczepankiewicz, F., Mårtensson, J., van Westen, D., Hansson, O., Westin, C.F., Nilsson, M., 2020. Towards unconstrained compartment modeling in white matter using diffusion-relaxation mri with tensor-valued diffusion encoding. Magnetic Resonance in Medicine 84, 1605–1623.

Lampinen, B., Szczepankiewicz, F., Mårtensson, J., van Westen, D., Sundgren, P.C., Nilsson, M., 2017. Neurite density imaging versus imaging of microscopic anisotropy in diffusion mri: A model comparison using spherical tensor encoding. Neuroimage 147, 517–531.

Lampinen, B., Szczepankiewicz, F., Novén, M., van Westen, D., Hansson, O., Englund, E., Mårtensson, J., Westin, C.F., Nilsson, M., 2019. Searching for the neurite density with diffusion mri: challenges for biophysical modeling. Human brain mapping 40, 2529–2545.

Le Bihan, D., Breton, E., Lallemand, D., Aubin, M., Vignaud, J., Laval-Jeantet, M., 1988. Separation of diffusion and perfusion in intravoxel incoherent motion mr imaging. Radiology 168, 497–505.

Mardia, K.V., Jupp, P.E., 1990. Directional statistics. volume 494. John Wiley & Sons.

Mueller, S.G., Weiner, M.W., Thal, L.J., Petersen, R.C., Jack, C., Jagust, W., Trojanowski, J.Q., Toga, A.W., Beckett, L., 2005. The alzheimer’s disease neuroimaging initiative. Neuroimaging Clinics 15, 869–877.

Novikov, D.S., Fieremans, E., Jespersen, S.N., Kiselev, V.G., 2019. Quantifying brain microstructure with diffusion mri: Theory and parameter estimation. NMR in Biomedicine 32, e3998.

Panagiotaki, E., Walker-Samuel, S., Siow, B., Johnson, S.P., Rajkumar, V., Pedley, R.B., Lythgoe, M.F., Alexander, D.C., 2014. Noninvasive quantification of solid tumor microstructure using verdict mri. Cancer research 74, 1902–1912.

Parker, D., Ould Ismail, A.A., Wolf, R., Brem, S., Alexander, S., Hodges, W., Pasternak, O., Caruyer, E., Verma, R., 2020. Freewater estimator using interpolated initialization (fernet): Characterizing peritumoral edema using clinically feasible diffusion mri data. Plos one 15, e0233645.

Plewes, D.B., 1994. Contrast mechanisms in spin-echo mr imaging. Radiographics 14, 1389–1404.

Scherrer, B., Schwartzman, A., Taquet, M., Sahin, M., Prabhu, S.P., Warfield, S.K., 2016. Characterizing brain tissue by assessment of the distribution of anisotropic microstructural environments in diffusion-compartment imaging (diamond). Magnetic resonance in medicine 76, 963–977.

Scherrer, B., Warfield, S.K., 2010. Why multiple b-values are required for multi-tensor models. evaluation with a constrained log-euclidean model, in: 2010 IEEE International Symposium on Biomedical Imaging: From Nano to Macro, IEEE. pp. 1389–1392.

Sotiropoulos, S.N., Jbabdi, S., Xu, J., Andersson, J.L., Moeller, S., Auerbach, E.J., Glasser, M.F., Hernandez, M., Sapiro, G., Jenkinson, M., et al., 2013. Advances in diffusion mri acquisition and processing in the human connectome project. Neuroimage 80, 125–143.

Stanisz, G.J., Wright, G.A., Henkelman, R.M., Szafer, A., 1997. An analytical model of restricted diffusion in bovine optic nerve. Magnetic Resonance in Medicine 37, 103–111.

Sudlow, C., Gallacher, J., Allen, N., Beral, V., Burton, P., Danesh, J., Downey, P., Elliott, P., Green, J., Landray, M., et al., 2015. Uk biobank: an open access resource for identifying the causes of a wide range of complex diseases of middle and old age. Plos med 12, e1001779.

Suzuki, H., Gao, H., Bai, W., Evangelou, E., Glocker, B., O’Regan, D.P., Elliott, P., Matthews, P.M., 2017. Abnormal brain white matter microstructure is associated with both pre-hypertension and hypertension. PLoS One 12, e0187600.

Szafer, A., Zhong, J., Anderson, A.W., Gore, J.C., 1995a. Diffusion-weighted imaging in tissues: theoretical models. NMR in Biomedicine 8, 289–296. doi:10.1002/nbm.1940080704.

Szafer, A., Zhong, J., Gore, J.C., 1995b. Theoretical model for water diffusion in tissues. Magnetic resonance in medicine 33, 697–712.

Tariq, M., Schneider, T., Alexander, D.C., Wheeler-Kingshott, C.A.G., Zhang, H., 2016. Bingham–noddi: mapping anisotropic orientation dispersion of neurites using diffusion mri. NeuroImage 133, 207–223.

Tournier, J.D., Calamante, F., Connelly, A., 2007. Robust determination of the fibre orientation distribution in diffusion mri: non-negativity constrained super-resolved spherical deconvolution. Neuroimage 35, 1459–1472.

Tournier, J.D., Smith, R., Raffelt, D., Tabbara, R., Dhollander, T., Pietsch, M., Christiaens, D., Jeurissen, B., Yeh, C.H., Connelly, A., 2019. Mrtrix3: A fast, flexible and open software framework for medical image processing and visualisation. NeuroImage 202, 116137.

Van Essen, D.C., Ugurbil, K., Auerbach, E., Barch, D., Behrens, T., Bucholz, R., Chang, A., Chen, L., Corbetta, M., Curtiss, S.W., et al., 2012. The human connectome project: a data acquisition perspective. Neuroimage 62, 2222–2231.

Veraart, J., Novikov, D.S., Fieremans, E., 2018. Te dependent diffusion imaging (teddi) distinguishes between compartmental t2 relaxation times. NeuroImage 182, 360–369. URL: http://www.sciencedirect.com/science/article/pii/S1053811917307784, doi:10.1016/j.neuroimage.2017.09.030.

Vestergaard-Poulsen, P., Hansen, B., Østergaard, L., Jakobsen, R., 2007. Microstructural changes in ischemic cortical gray matter predicted by a model of diffusion-weighted mri. Journal of Magnetic Resonance Imaging: An Official Journal of the International Society for Magnetic Resonance in Medicine 26, 529–540.

Villalon-reina, J.E., Nir, T.M., Thomopoulos, S.I., Salminen, L.E., Fick, R., Frigo, M., Deriche, R., Thompson, P.M. ADNI, 2020. Tracking microstructural biomarkers of alzheimer’s disease via advanced multi-shell diffusion mri scalar measures, in: ISMRM, p. 2020.

Wilcoxon, F., 1945. Individual comparisons by ranking methods. Biometrics Bulletin 1, 80–83.

Zhang, H., Schneider, T., Wheeler-Kingshott, C.A., Alexander, D.C., 2012. Noddi: practical in vivo neurite orientation dispersion and density imaging of the human brain. Neuroimage 61, 1000–1016.

